# Democratised single cell proteomics uncovers tumor associated macrophage signature in CYLD cutaneous syndrome skin tumors

**DOI:** 10.64898/2026.02.06.704354

**Authors:** Joseph Inns, Andrew Michael Frey, Matthias Trost, Neil Rajan

## Abstract

Single-cell proteomics (SCP) reveals cellular heterogeneity and biological insights inaccessible to bulk analysis. Existing limitations are cost, sample loss during processing, and accessibility to state-of-the-art instrumentation. We describe a label free SCP methodology in human tissue, combining fluorescence activated cell sorting (FACS), oil-immersion cell handling, mass-spectrometry, and neural-network derived spectral libraries which address these issues. We tested this methodology in a skin tumor syndrome, CYLD cutaneous syndrome (CCS), assessing tumor heterogeneity. Using a Bruker timsTOF HT platform we quantified > 4000 proteins, averaging ∼700 per cell, through a cost-effective pipeline without specialised liquid handling infrastructure. By utilising pre-existing bioinformatic tools from the scRNA-seq field we implemented a robust analysis methodology, discriminating between macrophages, dendritic cells and tumor keratinocytes, in an unbiased analysis of 419 CCS tumor cells. We validated the biological accuracy of cell annotations by cross referencing with each cell’s FACS markers. Furthermore, we identified a novel CCS tumor associated macrophage population which carried a tumor microenvironment remodelling signature. Our findings demonstrate an accessible SCP technology capable of yielding novel biological discoveries in clinical tissue.

**Graphical abstract:** 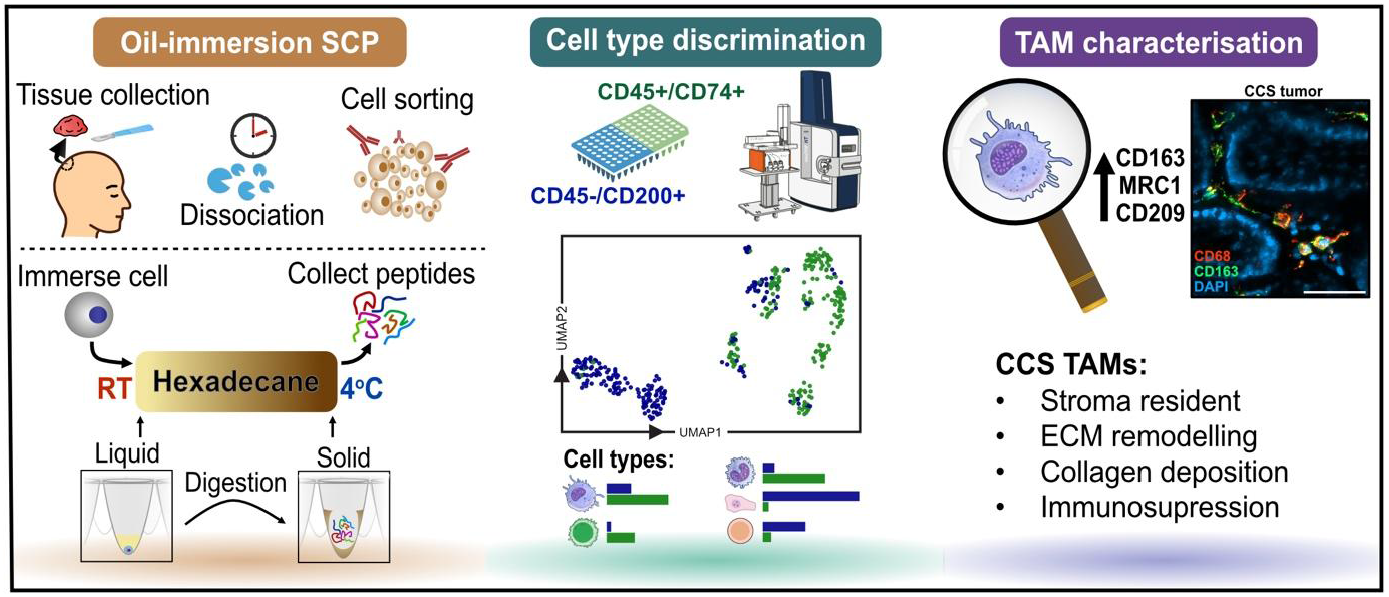

**IMPORTANT:**

- Manuscripts submitted to Review Commons are peer reviewed in a journal-agnostic way.
- Upon transfer of the peer reviewed preprint to a journal, the referee reports will be available in full to the handling editor.
- The identity of the referees will NOT be communicated to the authors unless the reviewers choose to sign their report.
- The identity of the referee will be confidentially disclosed to any affiliate journals to which the manuscript is transferred.

**GUIDELINES:**

- For reviewers: https://www.reviewcommons.org/reviewers
- For authors: https://www.reviewcommons.org/authors

**CONTACT:** The Review Commons office can be contacted directly at

## Introduction

Tumor heterogeneity determines therapeutic resistance^1^, metastatic potential^2^, biomarker interpretation^3^, tumor evolution^4^, and microenvironment interactions^5^. The development of novel therapeutic approaches is underpinned by the ability to delineate intra-tumor cellular heterogeneity, requiring methodologies with the resolution to elucidate these differences. Single cell technologies can reveal the diverse behaviour of cell types within biological tissues, enabling the identification of pathogenic traits which may go undetected in bulk analysis.

Single cell proteomics (SCP) is an emerging technology providing an unbiased characterisation of protein content at the cellular level. Despite its potential to reveal cell to cell heterogeneity and rare cellular states invisible to bulk analyses, widespread adoption of SCP remains constrained by its reliance on highly specialised instrumentation and expertise. Single cell RNA sequencing (scRNA-seq) represents an instructive precedent, having achieved widespread adoption through exponential cost reductions following Wright’s law^6^, and through the establishment of centralised facilities that democratised access to the technology. In contrast, advances in SCP have prioritised increasing proteomic depth and instrument sensitivity, frequently at the expense of sample throughput and cost^7,8^, limiting its accessibility to the broader research community. With recent advances enabling SCP to detect comparable numbers of unique proteins to transcripts detected by scRNA-seq, these complementary technologies are reaching functional parity in their molecular depth^9,10^. This convergence necessitates a strategic shift in SCP development priorities, from proteomic depth towards improved accessibility, throughput, and cost effectiveness to enable broader adoption of this powerful technology.

Here we describe an SCP methodology with a focus on reduced instrumentation requirements and decreased cost. We build upon the successful implementation of oil-immersion liquid handling described by others^11^, to capture single cells from enzymatically digested tumor tissue. We saw the potential of an accessible SCP workflow to study rare disease, where a clear pathogenesis and therapeutic target is often lacking. This led us to utilise our novel workflow to study CYLD cutaneous syndrome (CCS) skin tumors, arising in patients with pathogenic variants in *CYLD*. This tumor predisposition syndrome leads to the development and progressive growth of large hair follicle tumors which require repeated surgical removal to manage tumor burden^12^. We saw an unmet clinical need in characterising the cellular composition and pathogenic mechanisms driving tumor growth in CCS. SCP is uniquely positioned to address this challenge, providing direct quantification of long-lived proteins that define the pathogenicity of this tumor type and whose abundance is not directly related to transcriptional activity.

## Results

### Oil-immersion SCP improves protein detection and biological sensitivity

We sought to identify an accessible single cell proteomics methodology with the potential to reduce cost, improve user uptake, and accurately reflect biology. We utilised fluorescence activated cell sorting (FACS), to accurately dispense single cells, and allow enrichment of cell populations. We harnessed the increased throughput capabilities and reduced carryover of the Evosep liquid chromatography system coupled with a Bruker timsTOF HT mass spectrometer, representing an advanced yet economically practical instrument, where we have previously benchmarked low sample input^31^.

We prepared a patient tumor sample for SCP analysis by mechanical and enzymatic digestion followed by antibody labelling of a previously defined CD45^-^/CD200^+^ tumor keratinocyte population^13^ and a CD45^+^/CD74^+^ immune cell population. Single cells were FACS sorted (**Supplementary figure 1)** into hexadecane filled 96-well plates and incubated with a digestion mix, before loading onto Evosep tips for liquid chromatography-tandem mass spectrometry (LC-MS/MS) analysis (**Figure 1A**). The thermodynamic and hydrophobic properties of hexadecane allowed immersion of the single cell alongside the aqueous digestion mix solution at RT, whilst aqueous peptides could be retrieved by cooling and solidifying the hexadecane oil on ice followed by centrifugation (**Supplementary figure 2**). We first compared this oil-immersion methodology with an established C18 capture digestion method^14^ to assess performance over an indicative set of cells.

**Figure 1.**
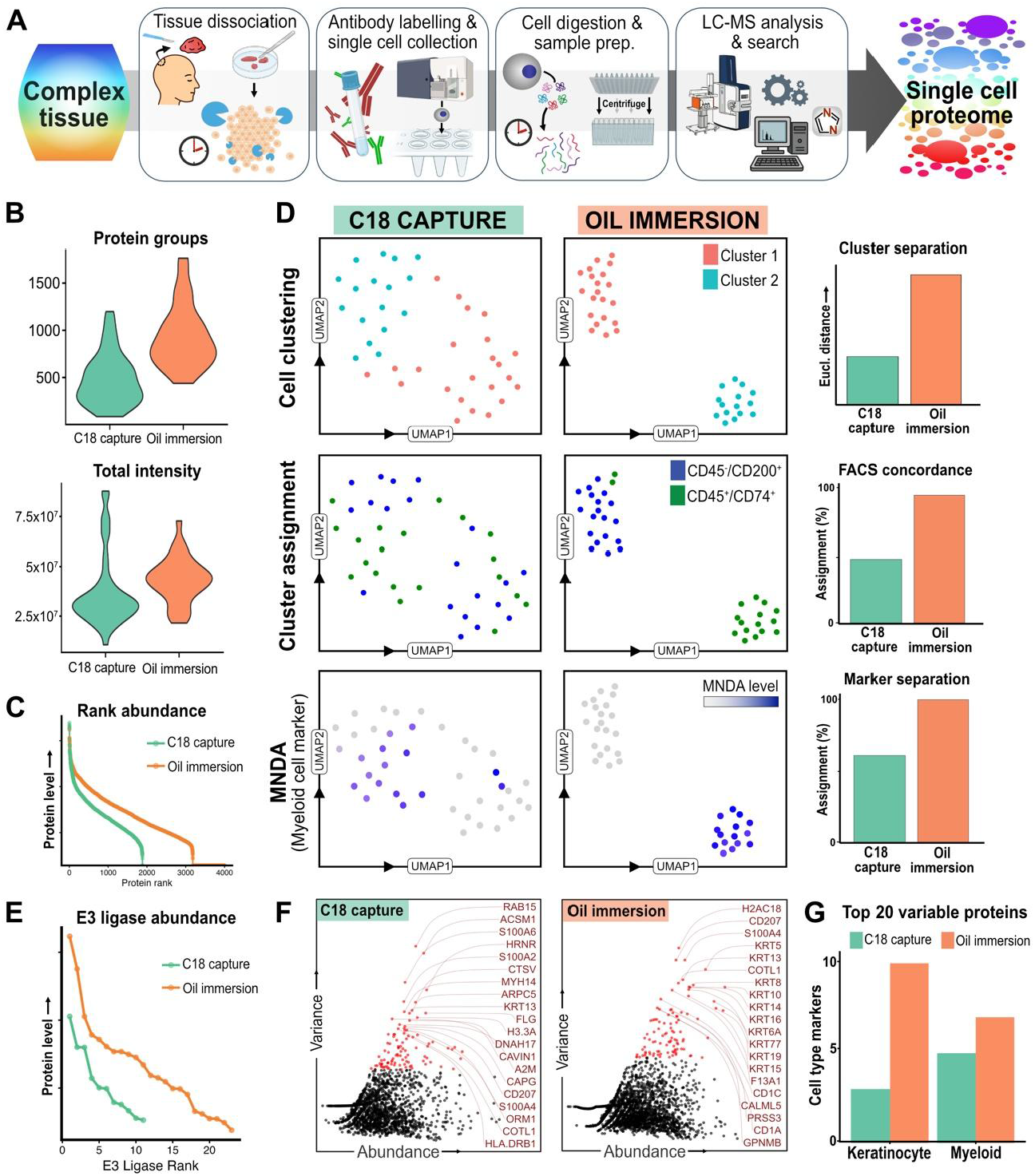
Oil-immersion single cell proteomics preserves characteristic biology. **A)** Workflow to generate single cell proteomics (SCP) data from a complex tissue using oil-immersion cell handling. **B)** Protein groups identified and total protein intensity per cell are compared between ‘C18 capture’ and ‘oil-immersion’ proteomics. **C)** Levels of proteins identified across SCP samples are ranked by abundance for C18 capture and oil-immersion methodologies. **D)** UMAP analysis of single cells from skin tumor tissue analysed by C18 capture or oil-immersion SCP workflows. Clustering distance (Euclidean), cell type clustering, and cell marker clustering are compared. **E)** Ubiquitin E3 ligases detected by SCP methodologies in rank order of intensity. **F)** The 20 most variable proteins from each SCP workflow were assessed **G)** and their cell type annotation over keratinocyte and myeloid cell types was plotted.

We identified increased protein intensity and protein group identifications per cell in the oil-immersion workflow (**Figure 1B**), as well as increased protein identifications across the single cells measured (**Figure 1C**). We also assessed each methodology in the following metrics: discriminating cells of different biological backgrounds (cluster separation), correctly assigning cells to their relevant population clusters (FACS concordance), and the presence of a myeloid specific marker (MNDA, marker separation) (**Figure 1D**). We found that the oil-immersion workflow outperformed the C18 capture workflow in each of these measures and correctly separated MNDA expressing cells from non-MNDA expressing cells on every occasion.

To understand the capability of each SCP workflow in identifying biologically relevant proteins we assessed the E3 ubiquitin ligases detected, which are present in biological systems across a large dynamic range. Eleven were detected in the C18 capture methodology and 23 in the oil-immersion methodology with an increased average expression, demonstrating increased biological sensitivity (**Figure 1E**). Furthermore, we investigated the most highly variable proteins in the combined CD45^-^/CD200^+^ and CD45^+^/CD74^+^ population, hypothesising these would be enriched for markers distinguishing each cell type (**Figure 1F**). Of the 20 most variable proteins, 10 of these were keratinocyte markers and 7 were myeloid lineage markers in the oil-immersion processed cells, whilst this dropped to 3 and 5 in the C18 capture workflow (**Figure 1G**). Therefore, we demonstrate by each metric that the oil-immersion workflow recognizes biologically relevant differences between cells as well as providing increased protein identifications and signal intensity.

### Oil-immersion SCP resolves biological heterogeneity

We performed a larger scale analysis of a CCS skin tumor using oil-immersion SCP, to explore the utility of this methodology in an increased throughput scenario. Each SCP run was searched against a library of 7872 proteins, created from ‘mini-bulk’ samples of 300 cells and predicted spectra. Using this library, the ‘mini-bulk’ proteomes covered 84.7% (6669 IDs) of this library, and the single cells covered 56.5% (4444 IDs) (**Figure 2A**). We also ran blank samples processed with digestion buffer but without a FACS dispensed cell. Here most blanks recorded zero protein identifications, both at the start and at the end of the sample set, highlighting low carryover and high detection specificity (**Figure 2B**).

**Figure 2.**
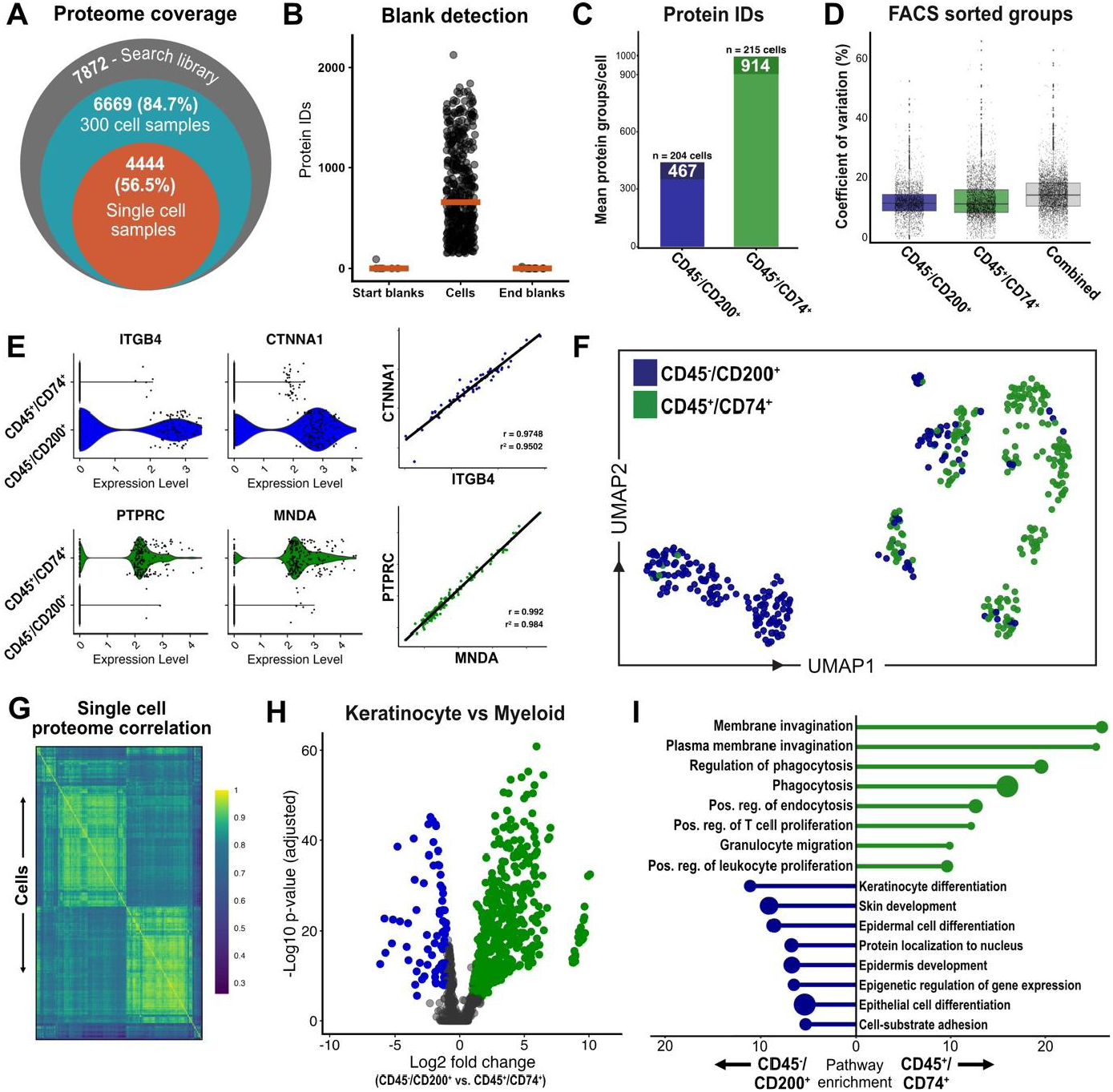
Oil-immersion SCP reveals CCS skin tumor heterogeneity. **A)** Protein identifications within the search library and samples acquired following the oil-immersion workflow: the combined 300 cell ‘mini-bulk’ samples and the single cell samples. **B)** Protein identifications per sample of start blanks which were run at the start of each 96-well plate set, end blanks ran at the end of each 96-well plate set, and single cells. **C)** Median number of protein identifications in CD45^-^/CD200^+^ and CD45^+^/CD74^+^ sorted cells. **D)** Assessment of workflow performance was made by coefficient of variation (%) per protein in FACS sorted cells, by sort group, and overall. **E)** ITGB4 and CTNNA1 levels were distinctly increased in CD45^-^/CD200^+^ tumor cells and their levels were highly correlated, while PTPRC and MNDA showed a similar pattern in CD45^+^/CD74^+^ cells. **F)** Uniform manifold projection (UMAP) of single cells from CYLD cutaneous syndrome skin tumors displays distinct separation based on FACS markers. **G)** Heatmap displaying the proteome level correlation of single cells in a CCS skin tumor. **H)** Differentially expressed proteins in CD45^-^/CD200^+^ and CD45^+^/CD74^+^ cells **I)** were enriched for biological processes that encompassed keratinocyte and immune cell processes. Bar length indicates fold enrichment, circle size indicates relative number of proteins per term.

After filtering and quality control we accepted 419 cells with a mean of 696 protein identifications per cell. The number of protein identifications differed in the FACS sorted cell groups, with a mean of 467 and 914 proteins per cell in CD45^-^/CD200^+^ and CD45^+^/CD74^+^ cells respectively (**Figure 2C**), representing biological differences between these two cell types. We also detected variation in the coefficient of variation (CV) per protein in each group (**Figure 2D**). The mean CV per protein was <20% and further decreased within sorted populations, indicating intra-group homogeneity. Population identities were confirmed by differential marker expression: CD45^-^/CD200^+^ cells showed elevated ITGB4 and CTNNA1 consistent with tumor keratinocytes, while CD45^+^/CD74^+^ cells exhibited increased MNDA and PTPRC consistent with myeloid cells (**Figure 2E**). Strong positive correlations between these marker pairs (r^2^ = 0.95 and 0.98, respectively) validated the biological fidelity of our proteomic measurements.

To assess whether the oil-immersion SCP workflow resolves distinct cell populations, we processed the data using Seurat^15^ and visualised cellular relationships via uniform manifold projection (UMAP) (**Figure 2F**). Here we observed well resolved cell clusters and importantly demonstrated separation between CD45^-^/CD200^+^ and CD45^+^/CD74^+^ sorted cells, without introducing a batch effect over separate 96-well plates (**Supplementary figure 3**). Additionally, hierarchical clustering and correlation analysis of individual single cells identified two majority clusters corresponding to the biological differences of the two FACS sorted cell types (**Figure 2G**).

Next, to identify biological differences between the CD45^-^/CD200^+^ and the CD45^+^/CD74^+^ sorted cells we performed differential protein expression analysis (**Figure 2H**) followed by gene set enrichment analysis of the differentially expressed proteins (**Figure 2I**). Here the enriched biological processes included keratinocyte differentiation and skin development in CD45^-^/CD200^+^ cells, and phagocytosis and granulocyte migration in CD45^+^/CD74^+^ sorted cells, matching the biology of their respective FACS sorted cell type. Collectively, these data confirm that oil-immersion SCP resolves meaningful cell type heterogeneity.

### Oil-immersion SCP enables unbiased cell type annotation and discovery

Our oil-immersion SCP methodology discriminated tumor cells based on their underlying biology, prompting us to explore what further biological insights could be drawn. We performed unbiased cell type annotation using SingleR^16^ and the human protein atlas database to further demonstrate the biological relevance of the data collected, and to allow downstream analysis of cell types of interest (**Figure 3A**). We found that 94% of the CD45^-^/CD200^+^ cells were annotated as keratinocytes, while 89% of the CD45^+^/CD74^+^ cells were annotated as either dendritic cells, monocytes or macrophages (**Figure 3B**). This analysis shows the utility of oil-immersion SCP in identifying heterogeneous cell types in complex clinical samples and led us to further explore the cell types in CCS tumors.

**Figure 3.**
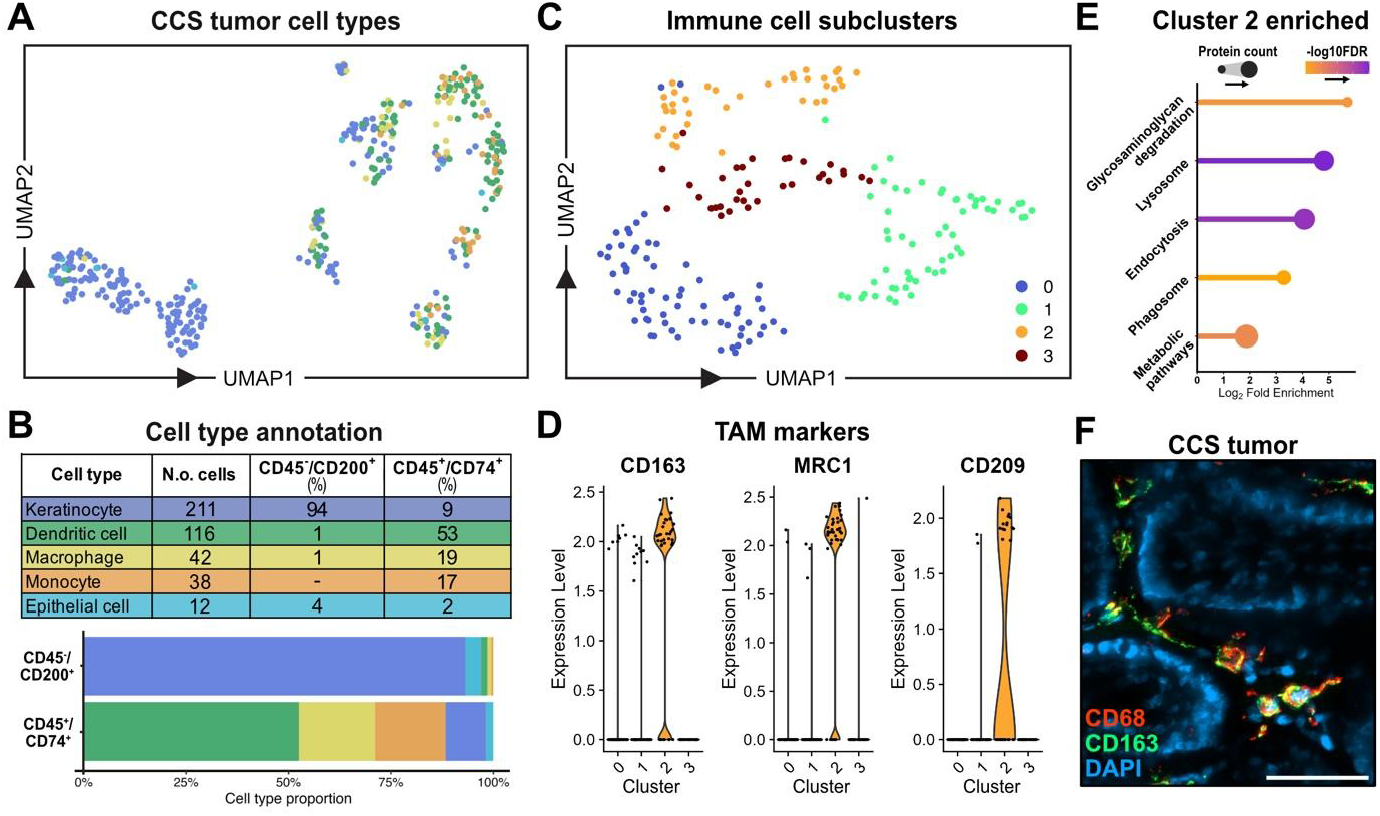
Oil-immersion based single cell proteomics identifies a tumor associated macrophage population. **A)** Unbiased cell type annotation of CCS tumor cells after oil-immersion based SCP analysis, and **B)** their cell type proportions. **C)** Re-clustering of CCS tumor immune cells to identify four immune cell subclusters. **D)** A CCS skin tumor subcluster displays tumor associated macrophage markers, **E)** and is enriched for KEGG pathway terms including lysosome and metabolic pathways. Bar length shows log2 fold enrichment of term, dot size shows number of proteins matched in the term, color displays decreasing p value. **F)** Immunofluorescence microscopy revealed presence of CD68/CD163 positive cells within CCS tumors. The scale bar is 50 μm.

We subclustered the annotated immune cells (dendritic cells, macrophages, and monocytes) identifying distinct clusters (**Figure 3C**). We probed the underlying biology of these clusters through differential protein expression analysis, where we observed 94 proteins that were markers of cluster 2 (Wilcoxon rank test, adj.p.val. < 0.05, log2FC > 1). These proteins included tumor associated macrophage (TAM) markers CD163, MRC1, and CD209 (**Figure 3D**). These markers are associated with M2 immunosuppressive macrophage populations and may explain the continued growth and immune surveillance escape of these tumors.

Gene set enrichment analysis revealed cluster 2 was enriched for KEGG pathway terms including lysosome, endocytosis and metabolic pathways (**Figure 3E**), suggesting this cell type is actively processing material to shape the tumor microenvironment (TME). This is further evidenced by enrichment of glycosaminoglycan degradation associated proteins, which play a prominent role in creating the TME architecture^17^. Furthermore, we investigated CCS tumor sections for the presence of cells expressing macrophage marker CD68, and TAM marker CD163. We observed CD68/CD163 positive cells interdigitating within CCS tumor stroma and between tumor ‘islands’ (**Figure 3F**). These findings further validate the utility of oil-immersion SCP and reveal novel CCS tumor biology which delineates tumor pathogenicity.

## Discussion

We demonstrate the application of a novel oil-immersion SCP workflow coupled with an informatic pipeline in a skin tumor sample and demonstrate its utility in delineating tumour cell heterogeneity and identification of cell populations which may facilitate tumorigenesis. Our accessible workflow demonstrates superior biological sensitivity and cell type resolution to an established approach^14^, without requiring specialised microfluidic instruments nor the latest generation mass spectrometer, facilitating broader adoption of SCP profiling. Analysis of 419 CCS tumor cells identified distinct keratinocyte and myeloid populations, including a population of macrophages expressing TAM markers, which we validated in tumor sections. Together these data show that an oil-immersion SCP approach is an effective methodology in identifying disease associated behaviours in heterogeneous tissues.

Unbiased proteomic profiling of single cells is a powerful tool to understand heterogeneous tissues but requires faithful representation of each cell to relay biological insight. Mass spectrometry based approaches have inherent biases for protein detection based upon peptide characteristics^18^, but equally importantly, sample processing steps can bias the resulting proteome^19^. For example, cell storage conditions can impact the resulting proteome^20,21^. Due to the large dynamic range in cellular protein abundance, reductions in protein recovery can also affect proteomics analysis, and workflows with multiple processing steps are more prone to reduced protein identification^22^. By encapsulating single cells in oil our workflow minimises processing steps and ensures reduced contact with plastics, a key determinant of peptide recovery^23^. Furthermore, we hypothesise that for institutions with access to more sensitive next generation mass spectrometers even greater biological depth and protein identifications will be possible whilst faithfully representing biology.

To investigate SCP data, we utilised existing bioinformatic tools developed in the transcriptomics field^15,16^. These tools provide a streamlined and standardised approach to single cell analysis but were developed with transcript data that does not necessarily correlate with protein abundance^24^. Consequently, cell annotation tools which are based on transcriptomic databases may not accurately label cell types when provided with their proteomic profile. Despite this limitation we evidence that these tools were concordant with our FACS sorting strategy and cell type marker proteins, indicating their utility but representing an area for refinement once larger SCP datasets become available.

Leveraging our unbiased cell type annotations, we subclustered myeloid cells to examine immune cell heterogeneity within CCS tumors and identified a distinct population of CD163^⁺^, MRC1^⁺^, and CD209^⁺^ cells, consistent with an alternatively activated (M2) macrophage phenotype^25^. This marker profile is characteristic of TAMs, specifically a mature, immunosuppressive subset shaped by TME signals^26^. These immunosuppressive M2 TAMs typically reside within the tumor stroma^27^, where they suppress T-cell responses and dampen anti-tumor immunity^28^. Beyond their immunomodulatory role, this population may also contribute to the sustained growth of CCS tumors through ECM remodelling. M2 TAMs are known to secrete matrix metalloproteinases as well as factors such as CCL18 that promote collagen deposition and ECM stabilisation^29,30^.

In summary, we demonstrate that oil-immersion SCP represents an accessible and biologically sensitive approach for interrogating heterogeneous clinical tissues. By minimising sample processing steps and reducing surface adsorption losses, our workflow achieved robust protein identification from individual cells isolated directly from a human skin tumor. The discovery of an immunosuppressive TAM population within CCS tumors was enabled by unbiased proteomic profiling, illustrating the capacity of this technology to reveal disease relevant biology obscured by bulk analyses. Importantly, this methodology does not require specialised microfluidics, extensive sample preparation, or the latest generation mass spectrometer, thereby lowering the barrier to entry for laboratories seeking to adopt SCP. As spectral library resources expand and annotation tools become tailored to proteomic data, we anticipate that accessible SCP workflows will become routine in clinical and translational research settings. The ability to resolve cellular phenotypes at the protein level, thereby identifying the functional effectors of cellular behaviour, positions SCP as a complementary and, in some contexts, superior modality to transcriptomic approaches for understanding tumor biology and identifying therapeutic vulnerabilities.

## Methods

### Patient samples

Research ethics committee approval was obtained from the Hartlepool research ethics committee and north east – Newcastle & North Tyneside research ethics committee (REC Ref:06/Q1001/59; 08/H0906/95+5;19/NE/0004). Skin tumor samples were obtained from patients with signed, informed, consent, with details in **Supplementary Table 1**.

### Single cell preparation

Patient tumor samples were dissociated with scalpels and digested with trypsin for 40 min at 37°C with agitation. Further digestion was performed with 1 mg/mL collagenase at 4°C and agitation overnight. A single cell suspension was obtained after passing through a 40 μm filter and the resulting suspension was labelled with CD45/CD74/CD200 antibody mix and DAPI (4′,6-diamidino-2-phenylindole) viability stain. Single cells were FACS sorted into wells of a 96-well plate containing 5 μL of solidified hexadecane and 1μL digestion buffer (0.2% DDM, 100 mM TEAB, 5ng/μL trypsin, 5ng/μL LysC). Plates were incubated at 45 °C for 2 h followed with quenching by addition of 30 μL 0.1% FA, 1% DMSO in water. Hexadecane was solidified on ice and plates were centrifuged at 500 g for 1 min at 4°C into prepared Evotips.

### Liquid chromatography mass spectrometry (LC-MS)

We analysed single cell samples with a Bruker timsTOF HT mass spectrometer in line with an EvoSep One liquid chromatography system (LC) operating the Whisper-Zoom 40 SPD method. The EvoSep LC injected samples onto a 15 cm Aurora Elite C18 column with integrated captive spray emitter (IonOpticks) at 50°C. Buffer A was 0.1% formic acid in HPLC water, buffer B was 0.1% formic acid in acetonitrile. The Bruker timsTOF HT MS was operated in diaPASEF (data-independent acquisition, parallel accumulation–serial fragmentation) mode with mass and ion mobility (IM) ranges of 300-1200 *m/z* and 0.6-1.45 1/K0. Four variable width IM-*m/z* windows each with two frames and no overlap were used as previously^31^ and provided (**Supplementary table 2**). TIMS ramp and accumulation times were 200 ms, total cycle time was ∼1.03 seconds. Collision energy was applied in a linear fashion, where ion mobility = 0.6-1.6 1/K0, and collision energy = 20 - 59 eV.

### Bioinformatic analysis

MS data files were searched in DIA-NN v2.0.1^32^ against a Uniprot *Homo sapiens* reference proteome (UP000005640) and a modified contaminant FASTA database^33^ at 1% FDR. MS1 and MS2 mass accuracy was set to 15 ppm, precursor range *m/z* 300-1200, and charge +2,+3,+4. A maximum of 2 missed cleavages and 2 variable modifications were allowed, with oxidation of methionine and N-terminal acetylation included as variable modifications. For the preliminary hexadecane versus C18 searches, no variable modifications and only 1 missed cleavage were allowed. The *in-silico* digest and spectral library prediction tool was used to search mini-bulk samples, and generate an empirical DIA-spectral library, which was then employed to search single cells. The output was processed in R using Seurat^15^ for normalization, UMAP creation and statistical tests. SingleR^34^ and the celldex R package utilising the Human protein atlas database were used for cell type annotation. Enrichment analysis was performed using ShinyGO 0.85^35^ and plots were generated with ggplot2^36^.

### Immunofluorescence microscopy

30 μm tissue sections were collected from snap frozen CCS tumors and fixed on glass slides in ice cold methanol for 10 min, followed by washing in Phosphate buffered saline (PBS). Sections were incubated in 30% sucrose in PBS w/v for 30 min and blocked in 0.5% bovine serum albumin (BSA), 0.3% Triton X-100 w/v in PBS for 1 h, RT. Incubation with anti-CD68 antibody (Cell signalling technology, 76437) was performed overnight at 4°C followed by washes in 0.2% BSA, 0.1% Triton X-100 w/v in PBS. Fluorescent staining was achieved with AlexaFluor 594 (Thermo Fisher Scientific) conjugated secondary antibody staining for 1 h at RT in the dark. After washes with 0.2% BSA, 0.1% Triton X-100 w/v in PBS, coverslips were mounted with ProLong Gold antifade with DAPI (Thermo Fisher Scientific). All washes were 3 x 10 min. Fluorescence images were captured with a Zeiss Axioimager Z2 microscope.

## Data availability

The mass spectrometry proteomics data have been deposited to the ProteomeXchange Consortium via the PRIDE partner repository^37^ with the dataset identifier PXD073250. Reviewers can access the datasets using the accession and this token: z9gGgCoCXcxl

## Acknowledgements

This research was partly funded by the NIHR Newcastle Biomedical Research Centre (BRC) awarded to the Newcastle upon Tyne Hospitals NHS Foundation Trust, Newcastle University and Cumbria, Northumberland, Tyne and Wear Foundation Trust.

## Supplementary Information

**Supplementary data 1 – Single cell proteome data (log2 normalised)**

**Supplementary data 2 – CCS tumor rds analysis object**

**Supplementary table 1.** Patient samples used in the study.

| ID | Age | Gender | Experiment | CYLD Genotype |
| --- | --- | --- | --- | --- |
| Sample 1 | 52 | F | Single cell proteomics (pilot) | c.2339delT |
| Sample 2 | 54 | F | Single cell proteomics | c.2460delC |

**Supplementary table 2.**
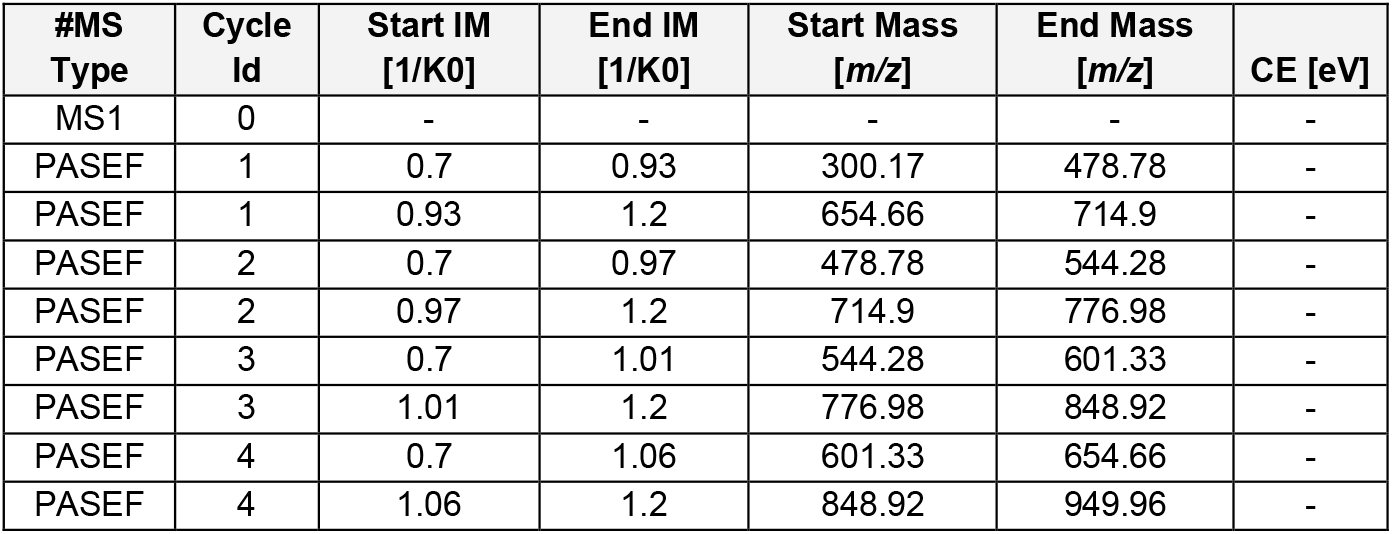
Bruker timsTOF HT diaPASEF acquisition scheme for single cell proteomics.

| #MS Type | Cycle Id | Start IM [1/K0] | End IM [1/K0] | Start Mass [m/z] | End Mass [m/z] | CE [eV] |
| --- | --- | --- | --- | --- | --- | --- |
| MS1 | 0 | - | - | - | - | - |
| PASEF | 1 | 0.7 | 0.93 | 300.17 | 478.78 | - |
| PASEF | 1 | 0.93 | 1.2 | 654.66 | 714.9 | - |
| PASEF | 2 | 0.7 | 0.97 | 478.78 | 544.28 | - |
| PASEF | 2 | 0.97 | 1.2 | 714.9 | 776.98 | - |
| PASEF | 3 | 0.7 | 1.01 | 544.28 | 601.33 | - |
| PASEF | 3 | 1.01 | 1.2 | 776.98 | 848.92 | - |
| PASEF | 4 | 0.7 | 1.06 | 601.33 | 654.66 | - |
| PASEF | 4 | 1.06 | 1.2 | 848.92 | 949.96 | - |

**Supplementary figure 1.**
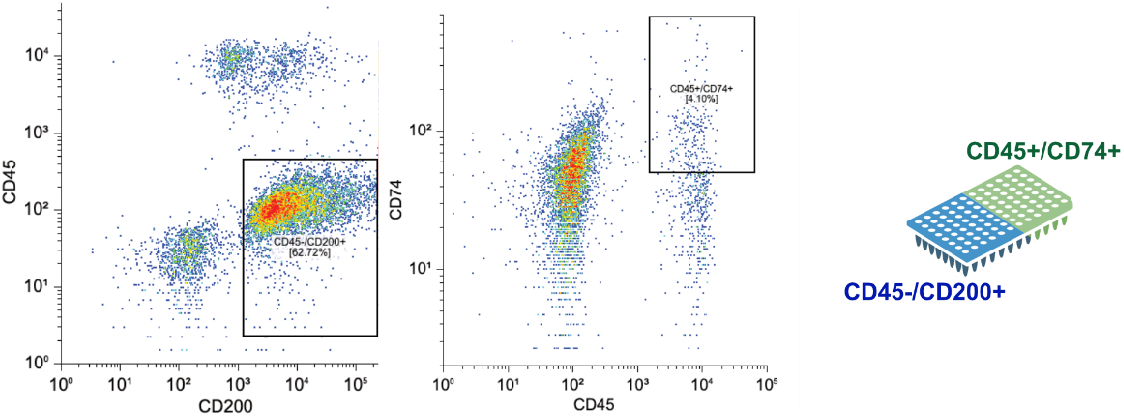
Fluorescence activated cell sorting (FACS) gating stratergy for the collection of CD45-/CD200+ tumor keratinocytes and CD45+/CD74+ tumor myeloid cells into 96-well plates.

**Supplementary figure 2.**
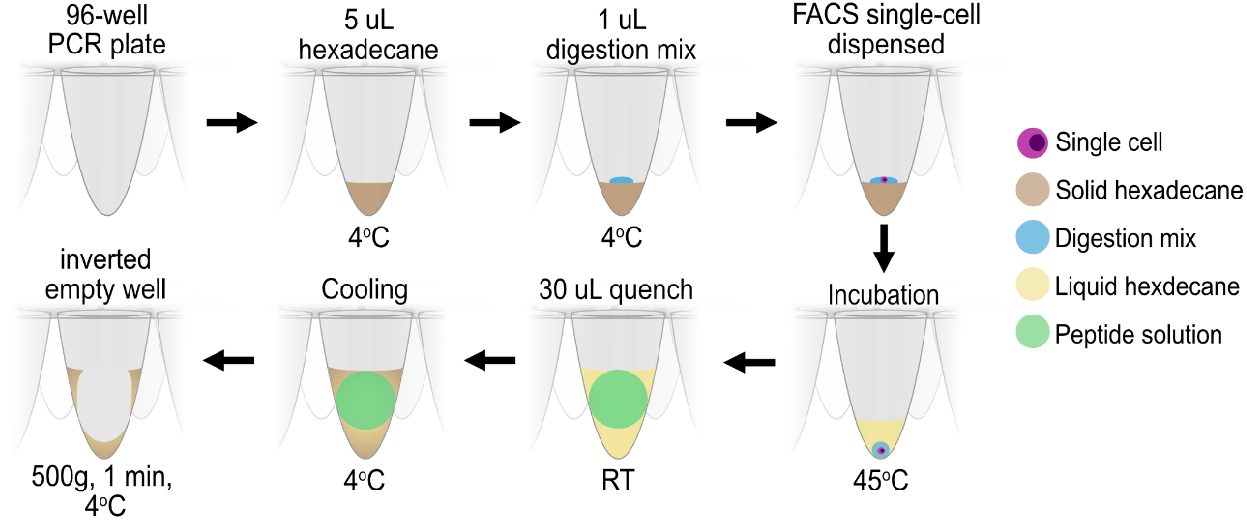
Single-cell handling workflow. Single cells were dispensed into wells of 96-well plates prepared with solidified hexadecane and a digestion mix. After fluorescence activated cell sorting of single-cells into each well, the 96 well plates were incubated at 45 °C, submerging the cell and digestion mix into the liquid hexadecane. After digestion was completed the reaction is quenched, the solutions cooled on ice and the aqueos peptide solution is removed by centrifuging the inverted plate.

**Supplementary figure 3.**
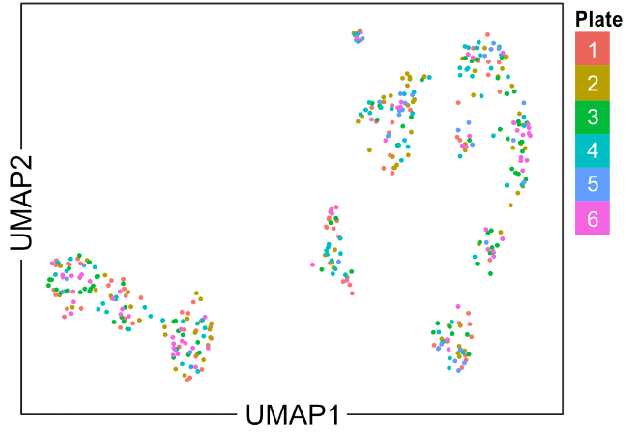
Uniform manifold projection of oil-immersion single cell proteomics analysed skin tumor cells. Six seperate plates of cells were collected and analysed and are distributed thoughout the UMAP.

